# Global patterns of plant form and function are strongly determined by evolutionary relationships

**DOI:** 10.1101/2023.01.13.523963

**Authors:** Pol Capdevila, Tom W. N. Walker, Franziska Schrodt, Roberto C. Rodriguez Caro, Roberto Salguero-Gomez

## Abstract

Plants display an incredible variety of forms and functions, which sustains a vast diversity of ecosystems on Earth. This diversity is the result of two forces: environmental filtering, the selection of phenotypes fit to a specific habitat; and evolutionary history, genetic constraints that determine the potential adaptations that species may evolve. To date, most studies have focused on describing macroecological patterns of trait variation, while the role of evolutionary history in determining these patterns has remained much less explored. Here, we combine a traits dataset including 4,213 vascular plant species with a phylogeny of the plant kingdom, to determine the influence of evolutionary history on the patterns of trait variation. Our results show that commonly used plant traits, such as plant height or leaf nitrogen, are strongly constrained by evolutionary history. Accounting for phylogenetic relationships changes the variance explained by the main axes of trait variation in the Plant Kindgom —the leaf economic spectrum and the plant size and organ axis continuum— with the former explaining more variance than the latter. Moreover, we demonstrate that the plant size continuum is more influenced by evolutionary history than the leaf economics spectrum. These results highlight the influence of evolutionary history in shaping plant life history strategies and the importance of accounting for phylogeny in trait-based studies.

## Main

Two main forces shape the combinations of traits found across the Tree of Life, namely evolutionary history and environmental filtering^1,2^. On the one hand, shared evolutionary history among descendants constrains the range of trait values that an organism can inherit^2^. On the other hand, organismal traits evolve to optimise the fitness of the individual according to its surrounding environment, *i*.*e*. environmental filtering^3–5^. Understanding the relative importance of these two processes is crucial to predict how traits assemble into successful life history strategies^1,6^, as well as to improve predictions on community assembly^7^ and ecosystem functioning^8^.

In recent decades, plant trait-based research has been fuelled by a multitude of comparative studies examining the main dimensions of trait variation from regional^9,10^ to global scales^6,11–13^. From these studies, general patterns have emerged, such as the spectra of leaf^14,15^, wood^16^, stem^17^, root^18^ as well as of form and function^6,11,13^. These spectra are underpinned through trade-offs between resource acquisition and allocation^1,19^. However, while analyses of global patterns of plant form and function sometimes do consider the degree to which trait variation is explained by environmental filtering - at least at a coarse level (e.g.^3^), the role of phylogenetic relationships in structuring these axes remains surprisingly unexplored (but see^20,21^). This consideration is important because evolutionary history can have a strong role in shaping trait variation^21,22^. Thus, accounting for phylogenetic relationships may alter our understanding of global patterns of trait variation^23^, and what these widely used dimensions are used for (*e*.*g*., quantify and predict ecosystem function).

Here, we explicitly quantify the role of evolutionary history in determining patterns of variation of plant traits at a macroecological scale. Specifically, we test the influence of evolutionary history on the established global spectrum of plant traits^6^. This spectrum postulates that most variability in aboveground plant traits can be captured by two axes: one related to plant size and their aboveground organs, and the other representing the leaf economics spectrum, representing the construction costs and quality of photosynthetic leaf area^6^. To do so, we couple a time-calibrated phylogeny^25^ for 31,749 seeding plants with trait information for 4,213 plant species from the TRY database^24^. We use the same traits as described in Díaz et al.^6^: plant height, leaf area, specific leaf area, leaf nitrogen concentration, specific stem density and seed mass. We hypothesise that: (H1) closely related species possess more similar trait composition than distantly related species (*e*.*g*., ^23,26^).; and (H2) that accounting for phylogenetic relationships among species will alter the global spectra of plant form and function.

### Traits have a strong phylogenetic signal across the plant kingdom

We quantified the phylogenetic signal of the six traits that define the plant form and function spectra to test the importance of evolutionary history. If traits show a strong phylogenetic signal, their variation among plant species is highly determined by the shared evolutionary history among lineages. This point is relevant because, if true, the assumption of independence of trait variation among species, often overlooked in macroecological studies of plant trait variation (*e*.*g*., ^6^), would be violated^27^. To evaluate the importance of the phylogeny on trait variation, we calculated the phylogenetic signal using Pagel ‘s *λ*^28^, with values close to 0 indicating a weak phylogenetic signal and values close to 1 indicating that trait variation is largely explained by evolutionary history^28^. We find that the six traits commonly used in the vast majority of plant trait studies research functional ecology are strongly influenced by evolutionary history across the plant kingdom (Table 1; Fig. 1). Contrary to previous suggestions that functional trait distinctiveness differs between lineages depending on the traits considered^29–31^, our analyses reveal that all six traits have a strong phylogenetic signal (*λ*>0.85; Table 1). Leaf area and specific leaf area are the two traits showing the strongest phylogenetic signal (*λ=*0.97), while plant height (*λ=*0.87) and seed mass (*λ=*0.90) showed the lowest values (Table 1). Because some traits were imputed for some species (see Methods), we repeated these analyses using only complete trait/species records. This validation exercise showed that the phylogenetic signal remained consistently high for all but one trait: seed mass. We attribute this difference in estimates of phylogenetic signal to the gap filling algorithm, which uses the taxonomic signal from correlations among other traits to also gap fill seed mass - although it is important to note that this inflation does not influence the identification of plant trait trade-offs we report here.

**Table 1.**
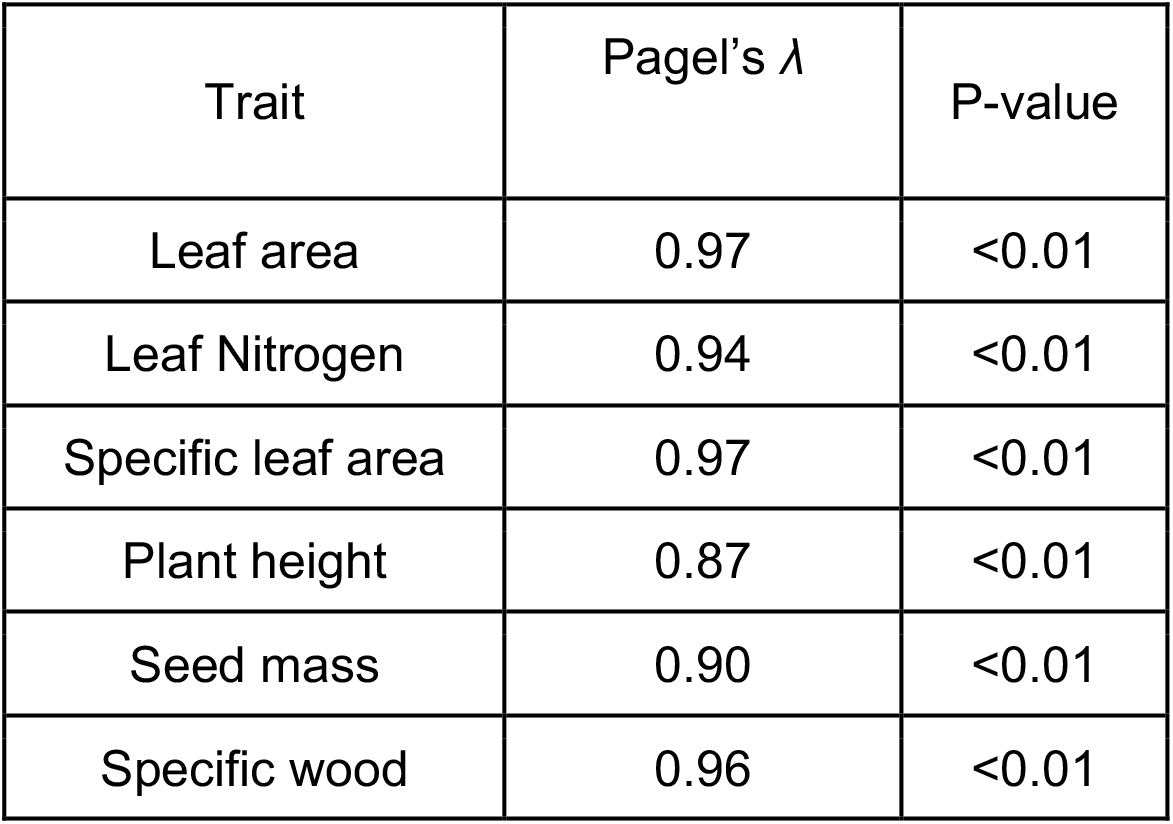
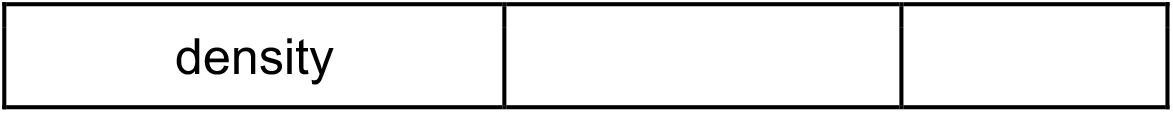
Six plant traits commonly used in trait-based research are strongly determined by phylogenetic relationships among species. Estimates of Pagel ‘s *λ* (± 95% C.I.) for six commonly used plant traits among 4,213 globally distributed vascular species. Values of λ close to 1 correspond to traits whose variation is strongly constrained by evolutionary history. The analyses presented here are for data imputed using a BHPMF approach (See Extended Methods), but are consistent with those emerging from non-imputed data.

| Trait | Pagel's $\lambda$ | P-value |
| --- | --- | --- |
| Leaf area | 0.97 | <0.01 |
| Leaf Nitrogen | 0.94 | <0.01 |
| Specific leaf area | 0.97 | <0.01 |
| Plant height | 0.87 | <0.01 |
| Seed mass | 0.90 | <0.01 |
| Specific wood | 0.96 | <0.01 |

**Fig. 1.**
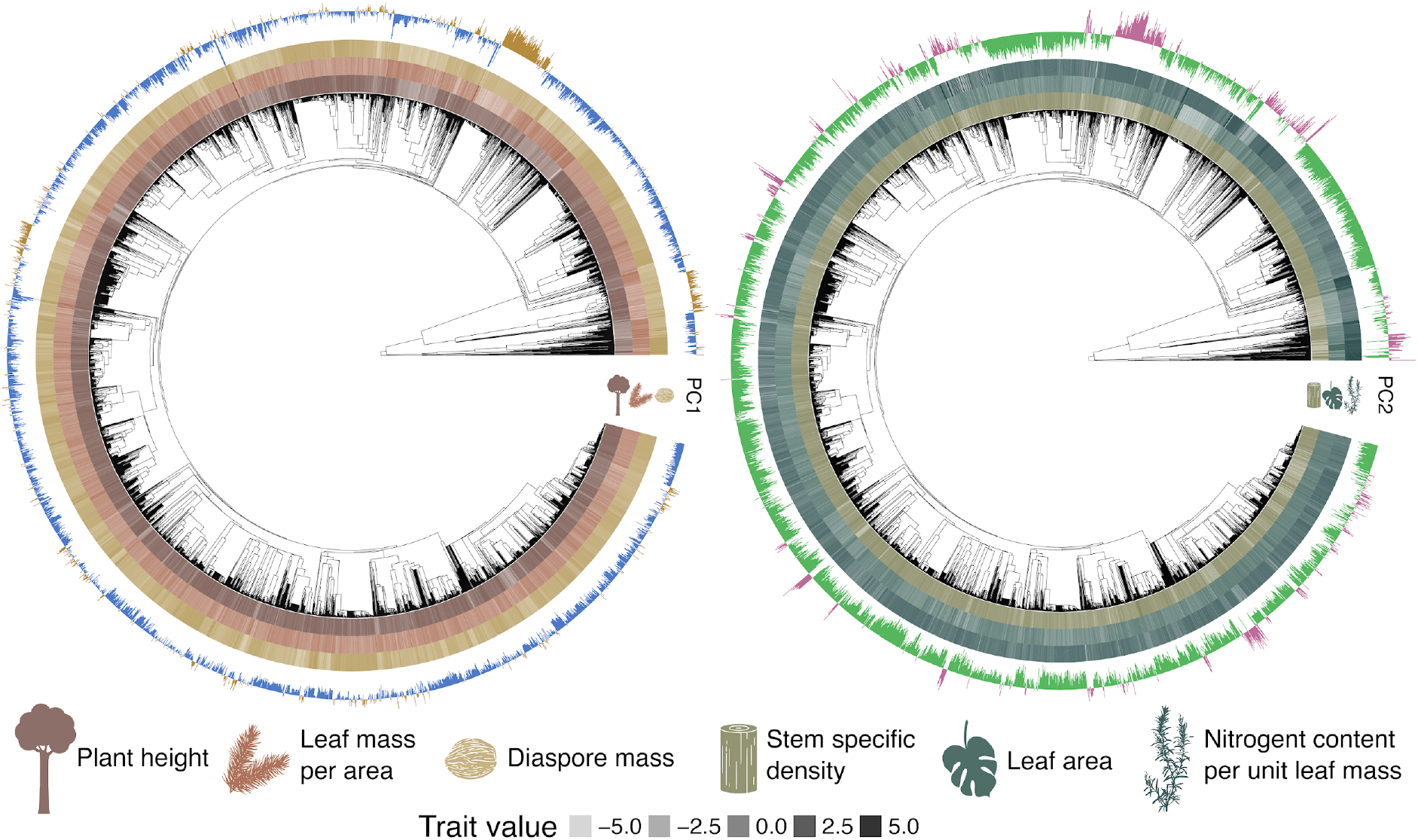
Evolutionary history has a strong influence on trait variation and resulting trait trade-offs across the plant kingdom. Traits and principal component scores displayed across the plant phylogeny of 4,213 plant species. (**a**) Phylogenetic trait variation (outer rings) that determine the first axis of variation (PC1; outermost ring) of plant form and function (plant height, leaf mass per area and diaspore mass). The first axis represents a trade-off between plant size and the development of their aboveground organs. (**b**) Phylogenetic trait variation (outer rings) that determine the second axis of variation (PC2; outermost ring) of plant form and function (stem specific density, leaf area and nutrient content per unit leaf mass). The second axis represents the leaf economics spectrum - a trade-off between the costs of leaf construction and growth.

### Evolutionary history influences the relative strength of plant trait trade-offs

To test whether the lack of evolutionary independence between species ‘ traits affects the interpretation of well-established dimensions of plant trait variation, we compared the results of two principal component analyses (PCA): with and without explicitly accounting for phylogenetic relationships. For the PCA corrected by phylogeny^27^ (pPCA), the composition of the main two trait axes are consistent with well-established trade-offs in plant form and function^6^. Principal Component 1 (PC1) absorbs30.82% of trait variance and describes the leaf economics spectrum^14^, *i*.*e*. a trade-off between large productive leaves that are poorly protected *vs*. small conservative leaves containing a high proportion of defensive compounds^6^. PC2 explains 22.13% of trait variance and represents a continuum of plant size, with tall, mostly woody, species with large leaves and large seeds at one end *vs*. short, mostly herbaceous, species with small leaves and small seeds at the other end^6^. Overall, the phylogenetically corrected PCA (pPCA) displays a strong phylogenetic signal, with a Pagel ‘s *λ* value of 0.96 (Fig. 2**a**). However, these two axes of trait variability have the same overall loading of traits as those identified by the non-phylogenetically corrected PCA (Fig. 2**b**). Specifically, plant height and seed mass continue to load similarly on the plant size continuum, while stem density continues to load oppositely to SLA and leaf nitrogen on the leaf economics spectrum (Fig. 2**a**,**b**). The relative positions of species with respect to both axes remain consistent irrespective of phylogenetic correction (Fig 2**c**), in that Euclidean distances between corresponding plant size and leaf economics PC scores of the two PCAs n are lower (mean: 1.41 ± 0.35 S.D.) than expected by chance alone (2.58 ± 1.44; Fig. 2**c**). These results collectively demonstrate that the main axes of trait variation can be appropriately identified, irrespective of whether evolutionary history of species is accounted for or not.

**Fig. 2.**
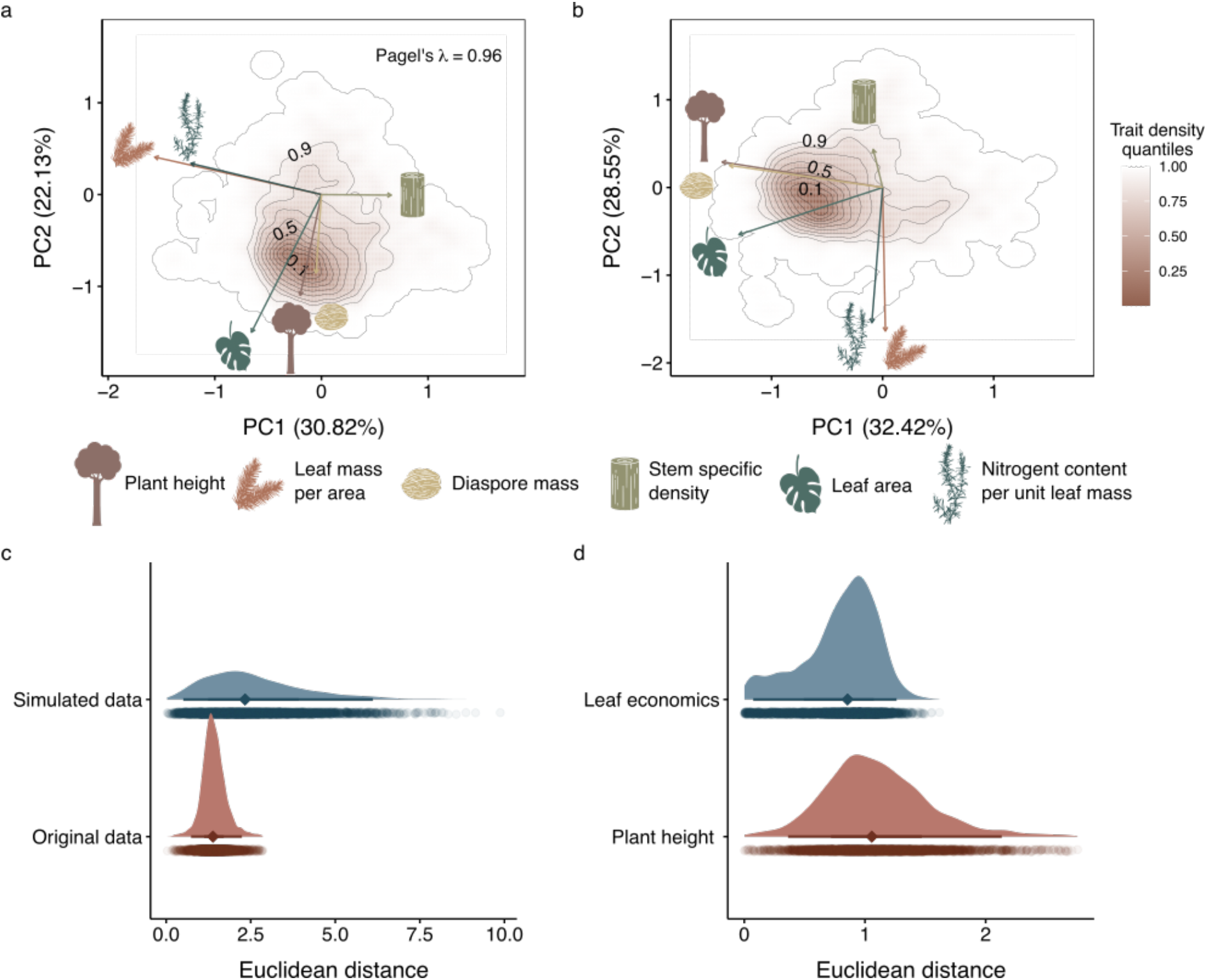
Plant form and function is constrained by species ‘ phylogenetic relationships. **a**. Phylogenetically-corrected Principal Component Analysis (pPCA) of six commonly used traits (See Table 1) across 4,213 plant species. The dominant axis corresponds to a continuum of organ size (leaf area, seed mass) and plant size (height), whereas the secondary axis captures the leaf economics spectrum (leaf mass area, leaf nitrogen, stem specific density). **b**. PCA of the same traits and data where phylogenetic relationships have not been considered. The relative importance of the leaf economics spectrum when correcting by phylogeny becomes greater (pPC1, 30.82%) than in the PCA (PC2 28.55%). Likewise, the plant size continuum decreases its relative importance when comparing the PCA with (pPC2, 22.13%) and without phylogenetic correction (PC1, 32.42%). **c**. Distribution of Euclidean distances among the species positions between rotated axes of the pPCA *versus* PCA. The simulated data represent Euclidean distances among species in the pPCA and PCA, with the positions of species randomised in the pPCA. Euclidean distances in the original data are lower than for simulated data. **d**. Distribution of Euclidean distances of species positions between the pPCA and PCA for the plant size continuum and the leaf economics spectrum. Euclidean distances are larger for the plant size axis than the leaf economic spectrum, suggesting that the former is more constrained by evolutionary history.

Despite the compositional similarities between the pPCA and PCA, accounting for phylogeny does alter the relative importance of the two main axes of trait variation (Fig. 2). Specifically, for the pPCA, the leaf economics spectrum is more important (PC1) than the plant size continuum (PC2), with the inverse being true for the uncorrected PCA. These differences are also apparent in the total variance explained by PC1 and PC2, which increases from 52.95% (Fig. 2a) to 60.97% (Fig. 2b). Species-wide Euclidean distances among PCAs with and without phylogenetic correction are also higher for the plant size continuum axis (mean: 2.60 ± 1.44 S.D.) than the leaf economics spectrum axis (1.10 ± 0.43; Fig. 2c). However, the traits characterising the plant size continuum are not those that show the strongest phylogenetic signal overall (Table 1). These findings demonstrate that evolutionary history controls the plant size continuum more strongly than the leaf economics spectrum, and - once phylogeny has been accounted for - that the leaf economics spectrum describes considerably more variation in plant form and function.

## Discussion

We combine the most comprehensive aboveground trait database and plant kingdom phylogeny available to show that evolutionary history plays a key role in shaping aboveground trait variation across the plant kingdom. We not only find that plant traits have a strong phylogenetic signal, but also that accounting for the influence of evolutionary history changes the relative importance of well-established axes of trait variation to plants globally - *i*.*e*. the leaf economic spectrum and the plant size continuum. In short, these results highlight the importance of accounting for phylogeny in trait-based studies, at least at the macroecological scale.

Our findings are consistent with recent studies showing a strong influence of evolutionary history on traits^20,21,29,32^. Indeed, the existence of a strong phylogenetic trait signal is commonly exploited to gap fill trait databases (*e*.*g*., phylogenetic imputation^33^) and estimate phylogeny from spectral remote sensing (“spectral traits “)^34^. However, previous explorations of phylogenetic signals for plant traits have been isolated to specific taxonomic groups^21,29,32^, with few global exceptions^21^. Nevertheless, it is unsurprising that using a large compilation of trait data spanning 403 families and 7,079 genera of plants we find a strong phylogenetic signal. What is not possible to tease apart from our approach is whether these patterns are exclusively a result of evolutionary history or a product of eco-evolutionary processes, as previously hypothesized^20,35^. Moreover, It is worth noting that our study focuses on plant traits measured in ambient conditions, whereas phylogenetic signals have been suggested to be low when plants are exposed to extreme conditions, such as drought^36^ or warmth^21^. Regardless of this, our findings that phylogenetic trait signals are strong for every trait when using a large number of plant species strongly advocates for the incorporation of phylogenetic correction in global trait analyses. This is particularly important when considering that many statistical analyses assume independence among species traits.

The change in importance of the first two dimensions of trait variation when incorporating evolutionary history has important implications for our current understanding of plant form and function. First, the fact that the constitution of major axes of trait variation remains the same with and without phylogenetic correction confirms that both axes are the result of inescapable trade-offs that shape the evolution of plant life history strategies^6^. Second, the change in the importance of the main axes of trait variation reflects that these are subject to different selective pressures^27^, with the plant size continuum being more constrained by evolution than the leaf economics spectrum. A recent study found that plant size continuum related traits show stronger latitudinal patterns than leaf traits^13^, which the authors attributed to the fact that plant size traits were more influenced by climate related variables while leaf traits were more determined by soil conditions^13^. However, our results seem to indicate that eco-evolutionary process must play an important role in determining these patterns, particularly for plant size related traits.

Finally, our results have important methodological evidence for using phylogenetic correction in further analyses. We demonstrate that neglecting the shared ancestry of species from trait analyses can alter the emergence, and thus interpretation, of patterns of trait variation. Previous studies have also suggested that ignoring evolutionary history when traits have a phylogenetic signal can lead to statistical inaccuracies, such as a decrease in estimation accuracy or inflated type I error^26,37^. Therefore, analysing trait data without accounting for evolutionary history entails a substantial risk of misinterpreting general patterns of traits variation. While here the constitution of main trade-offs among traits remains unaltered when accounting for phylogeny, the fact that their relative importance changes begs for a reconsideration of existing life history theory and raises concerns about the validity of previous approaches that do not consider phylogenetic relationships. We thus encourage researchers to consider the potential risks of ignoring evolutionary interdependence among species and call for future efforts to improve the reliability and robustness of trait-based approaches using evolutionary history.

## Methods

### Trait data

We obtained plant traits data from the open-access version of the TRY database^24^ (version 5; downloaded in April 2019). The traits are: plant height (m), seed mass (mg), stem specific density (mg mm^-3^), leaf area (mm^2^), specific leaf area (mm^2^ mg^-1^) and leaf carbon (mg g^-1^). We selected these set of traits because of their key role in determining the plants form and function spectrum^6,38^. After removing clear outliers (reported error risk ≥ 4; values falling outside of documented limits), we calculated species-wise means and gap-filled missing trait values on the full dataset (N = 46905 species) using a two-step process. First, we performed a Bayesian hierarchical probabilistic matrix factorisation imputation (package “BHPMF”)^39^, which is a taxonomically-constrained gap-filling approach that has been previously applied to the TRY database^6^. The imputation was repeated 90 times, each time starting with a different combination of parameters (per-fold samples = 900-1000; cross-validation steps = 10-20; burn-in steps = 10% data length) and filtering out extreme (>1.5 times the maximum observed value for a trait) or highly uncertain (coefficient of variation >1) gap-filled values. We then calculated means of remaining gap-filled values across all iterations and used these to replace missing cases in original trait data. Second, we performed five iterations of multivariate imputation by chained equations (package “mice”^40^) on the partially gap-filled dataset and replaced remaining missing cases with the mean values from all iterations.

### Phylogenetic data

To account for the evolutionary history of the species in our analyses, we built a phylogeny using the *V. PhyloMaker* R package^41^. *V. PhyloMaker* allows building a rooted and time-calibrated phylogeny using a species list based on already built plant phylogenies^25,42^.

### Phylogenetic signal

To measure the influence of the evolutionary history on the patterns of trait variation across the different species of plants we used Pagel ‘s λ. Pagel ‘s λ quantifies the strength of the phylogenetic relationships on trait evolution under a Brownian motion model^2,43^. This metric varies between 0, when the observed patterns are not due to phylogenetic relationships, and 1 when the observed patterns can be explained by the employed phylogeny^2,44^.

### Multivariate analyses

To explore the main axes of plant traits variation and the influence of evolutionary history on them, we performed principal components analysis with and without phylogenetic correction (pPCA and PCA, respectively). PCA is a multivariate analysis that reduces a set of correlated variables into linearly uncorrelated measurements, the so-called principal components (PCs). To account for shared ancestry in the PCA we corrected it using the phylogeny (pPCA^27^). The pPCA considers the correlation matrix of species ‘ traits while accounting for phylogenetic relationships and simultaneously estimating Pagel ‘s λ with maximum likelihood methods. This approach allows us to accommodate residual errors according to a variance–covariance matrix that includes ancestral relationships between any pair of species from our phylogenetic tree^44,45^. The variance–covariance matrix represents the expected covariance between species ‘ trait values, given a phylogenetic tree and under a specific model of evolution^27^. The expected covariance between species ‘ trait values is directly proportional to the distance between the species and their most recent common ancestor, that is measured as the branch length of the phylogeny^27^. The pPCA was estimated using the *phyl*.*pca* function from the r package phytools^45^, assuming a Brownian motion model of evolution (Revell, 2010)^44^. Plant traits data were log- and z-transformed (M = 0, SD = 1) to fulfil normality assumptions of PCAs^46^. To test for the influence of the evolutionary history in the positioning of the species in the multivariate space, we calculated the Euclidean distances among the species position between the rotated axes of the pPCA and PCA. Then, we randomised the position of species in the pPCA and measured again the Euclidean distances among the species in pPCA and PCA. The comparison between the true distances and the ones obtained with the simulated data indicates whether including the evolutionary history modifies the position of species within the multidimensional space more than expected by chance. To explore the influence of evolutionary history on the two main axes of trait variation, we measured the Euclidean distances of the species position between the pPCA and the PCA for the plant size continuum and the leaf economic spectrum. The axis with the larger Euclidean distances would indicate a stronger influence of evolutionary history.

## Acknowledgements

This study was part funded by a Ramón Areces Foundation Postdoctoral Fellowship to PC (BEVP30P01A5816) hosted by RSG. RSG was supported by a NERC Independent Research fellowship (NE/M018458/1).

## Author contributions

Conceptualisation: PC, RSG, TWNW, and FS. Data collection: TW and FS. Data curation: PC, TWNW, and FS. Formal analysis: PC in close collaboration with RR and TWNW, and the support of RSG and FS. Writing: PC and RSG, with support of all authors. Reviewing and editing: all authors.

